# BODIPY-Tagged β-Lactams as Selective Quenched Activity-Based Probes to Target Human Neutrophil Elastase

**DOI:** 10.64898/2026.03.19.712884

**Authors:** Rita Felix, Luís Carvalho, Ismael Rufino, Rita Guedes, Ana Margarida Madureira, Ana Mallo-Abreu, Lídia Gonçalves, Olga Genilloud, Rosario Fernandez-Godino, Maria C Ramos, Rui Moreira

## Abstract

Activity-based probes are indispensable tools for interrogating protease function, and quenched fluorescent variants enable dynamic, real-time imaging of enzymatic activity. Despite these advances, very few quenched activity-based probes (qABPs) have been reported for serine proteases, which constitute the largest and most diverse mechanistic class of proteases. *β*-Lactams have been extensively used to develop molecular tools and drugs designed to bind or be hydrolysed by serine-dependent bacterial enzymes. Here, we report the first monocyclic *β*-lactam-containing qABPs for detecting serine proteases. Both the enzyme-triggered activation mechanism and intrinsic reactivity of the probes were highly dependent on the relative position of the BODIPY-FL fluorophore and quencher moiety at the *β*-lactam core. qABPs displaying the most efficient turn-on mechanism were shown to selectively target human neutrophil elastase (HNE) in different human cell lysates. The most successful qABP was rapidly internalised and targeted HNE in U937 cells and human neutrophils. These results demonstrate the potential of the modular *β*-lactam warhead to develop turn-on probes to track neutrophil serine proteases in live cells

## Introduction

Serine hydrolases encompass a large superfamily of enzymes that play key functions in biological processes in mammals, bacteria and viruses. Despite their importance, the role of many serine hydrolases in the development and progress of several diseases remains poorly characterized.^1-3^ Human Neutrophil Elastase (HNE) is a serine hydrolase of the chymotrypsin family expressed in polymorphonuclear neutrophils and has been associated with lung related diseases, such as cystic fibrosis, chronic obstructive pulmonary disease and acute respiratory distress syndrome.^4, 5^ It has also been recently reported that HNE is present in the tumour microenvironment, promoting tumour growth and metastasis.^6-8^ However, the impact of HNE released to the tumour microenvironment on cancer progression remains largely unexplored due to the lack of selective chemical tools to study enzyme localization and dynamics in cells and tissues.

Activity-based probes (ABPs) are small molecule tools used in chemical proteomics and cell imaging, incorporating an electrophilic group that covalently reacts with catalytic amino acid residues and a reporter tag for detection and enrichment of probe-labelled enzymes.^1, 9^ The selectivity displayed by ABPs is strongly dependent on the structural motifs required for optimal molecular recognition by the target enzyme and the intrinsic reactivity of the warhead.^9^ ABPs containing a fluorophore tag enable both sensitive gel-based visualization and fluorescence imaging of enzymatic activity, thus being extensively used as reagents for the detection and study enzymes in cells. However, background signal generated by intrinsic fluorescence of the probe prevents successful application in real-time imaging with ABPs. Achieving significant signal-to-background signal typically requires extensive washing steps *in vitro* or clearing times *in vivo* to reduce the background signal. A solution for this problem is offered by quenched ABPs (qABPs), in which a fluorescence quencher is incorporated as a leaving group. A requirement for this approach is the presence of a warhead with a single leaving group connected to a fluorescence quencher. A key advantage of qABPs over traditional ABPs is their fluorescence activation exclusively upon target enzyme binding, eliminating background signal from unreacted probe in solution and thereby overcoming major imaging limitations (Figure 1).^10-13^

**Figure 1.**
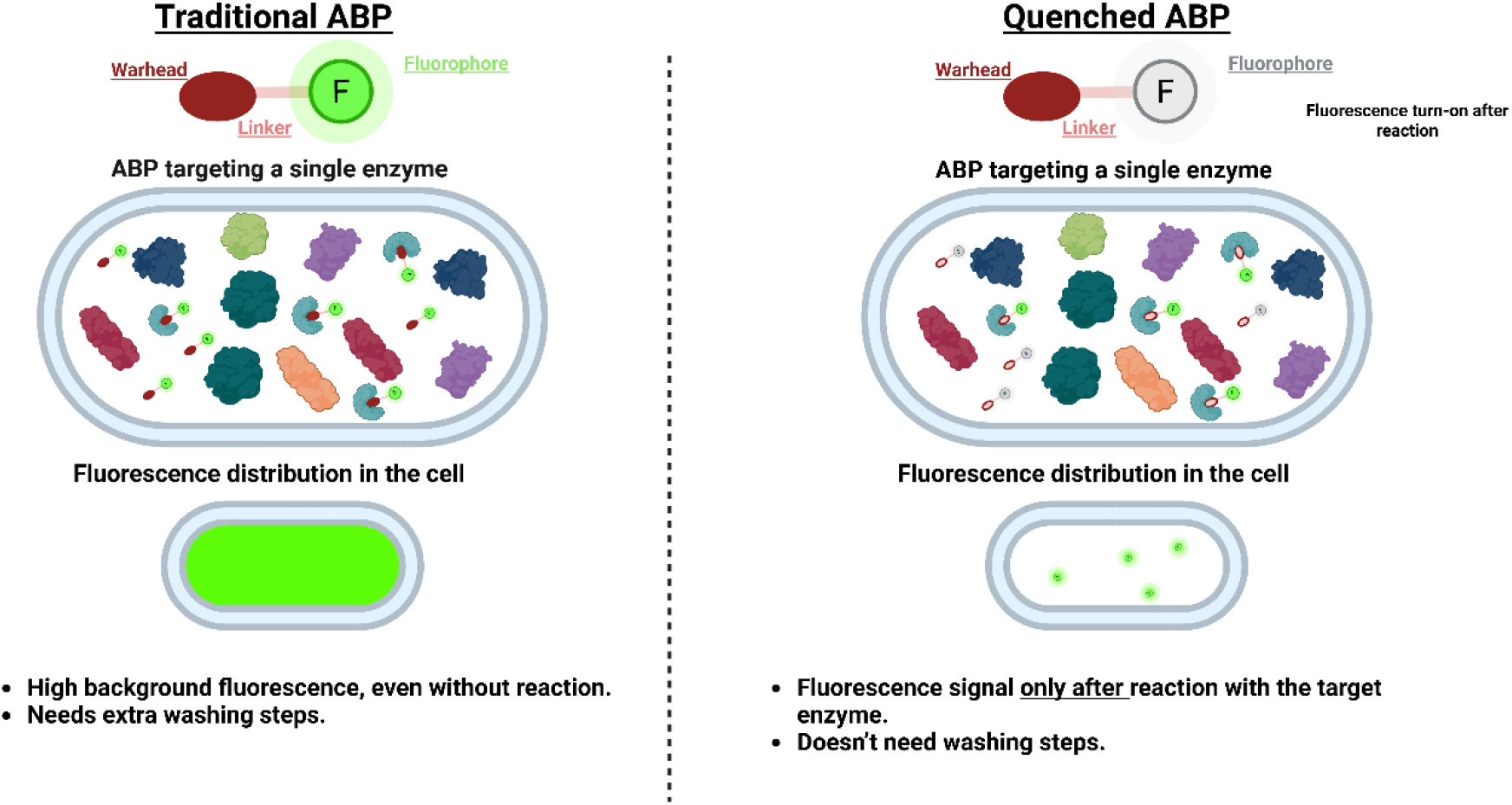
Activity-Based probes (ABPs) vs quenched Activity-Based Probes (qABPs) in biological applications.

While qABPs for cysteine proteases are well established experimental tools, their counterparts designed to target serine hydrolases, including HNE, remain largely underexplored.^14-23^ The first near-infrared fluorogenic off–on probe reported for HNE, incorporating a hemicyanine fluorophore, was shown to restore fluorescence upon reaction with HNE (Figure 2A).^24^ However, because the fluorophore is released into solution, this strategy is not well suited to gel-based assays or pull-down experiments. Alternatively, ABPs can incorporate a linker between the reactive group and the tag, which functions not only as a spacer but also as a key determinant of selectivity. A common approach in cysteine-targeted ABPP involves introducing a short amino-acid sequence to direct a broadly reactive chemotype toward a specific protein. Using this principle, Rios et al.^25^ developed “triple qABPs” containing three fluorescein moieties paired with three DABSYL quenchers via a peptide sequence, which were selectively cleaved by HNE between isoleucine and norleucine (Figure 2B). While multiple fluorophore/quencher pairs is a well-established strategy to reduce background and amplify signal, it substantially increases the overall molecular weight. More recently, cell-permeable qABPs based on the broadly reactive phosphinate ester warhead enabled finetuning of selectivity across different protein targets by varying the amino-acid sequence between the fluorophore and the linker (Figures 2C,D), underscoring the critical role of linker design in qABP performance.^26, 27^ Nevertheless, incorporating peptide-based linkers increases both structural complexity and synthetic demands.

**Figure 2.**
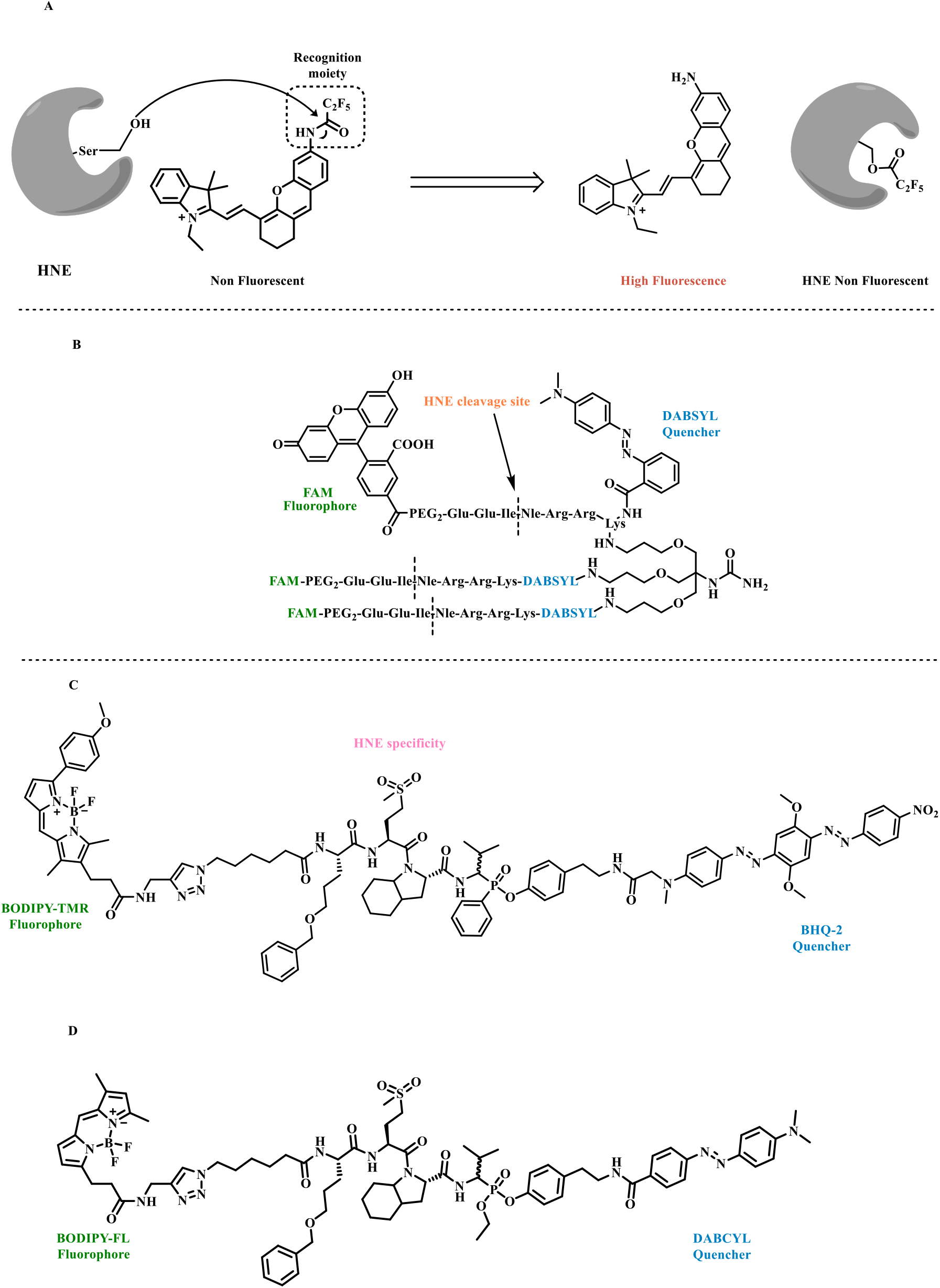
Examples of qABPs that target HNE. **A**. qABP where selective activation by HNE releases the amino xanthene fluorophore. **B**. Peptide-based “triple qABP”, comprising three fluorophore/quencher pairs connected through a peptide sequence selectively cleaved by HNE. **C/D**. Phosphonate ester qABPs, where the selectivity is determined by the amino acid sequence in the linker.

Monobactams are a class of narrow-spectrum *β*-lactam antibiotics with a unique monocyclic ring structure, that exert their activity by reacting with a conserved active-site serine residue in penicillin-binding proteins. While being used primarily to tackle infections caused by aerobic Gram-negative bacteria, monobactam moieties have also been employed to inhibit human serine hydrolases, including HNE ^28-31^ and Dipeptidyl Peptidases 8 and 9.^32^ Herein, we report the first qABPs featuring a monocyclic *β*-lactam warhead and endowed with a novel turn-on mechanism. These probes demonstrated strong inhibition and high selectivity toward HNE, enabling its detection in complex proteomes.

## Results and Discussion

### Design and chemistry

The design of the monobactam qABPs relies on a mechanism-based enzyme inhibition to enable the formation of a stable acyl-enzyme complex. Two types of qABPs are reported, differing in the positions where fluorophore and quencher are installed, as well as on their activation and quencher-releasing mechanism (Figure 3). While type I probes contain the quencher as part of a *N*-1 acyloxymethyl moiety, their type II counterparts present the quencher at C-4 of the 4-membered ring. In both cases, *β*-lactam ring-opening upon reaction with a catalytic serine residue can promote a 1,2-elimination to release the quencher and form an electrophile that can react with a second amino acid residue close to the active site, resulting in an irreversible inhibition mechanism.

**Figure 3.**
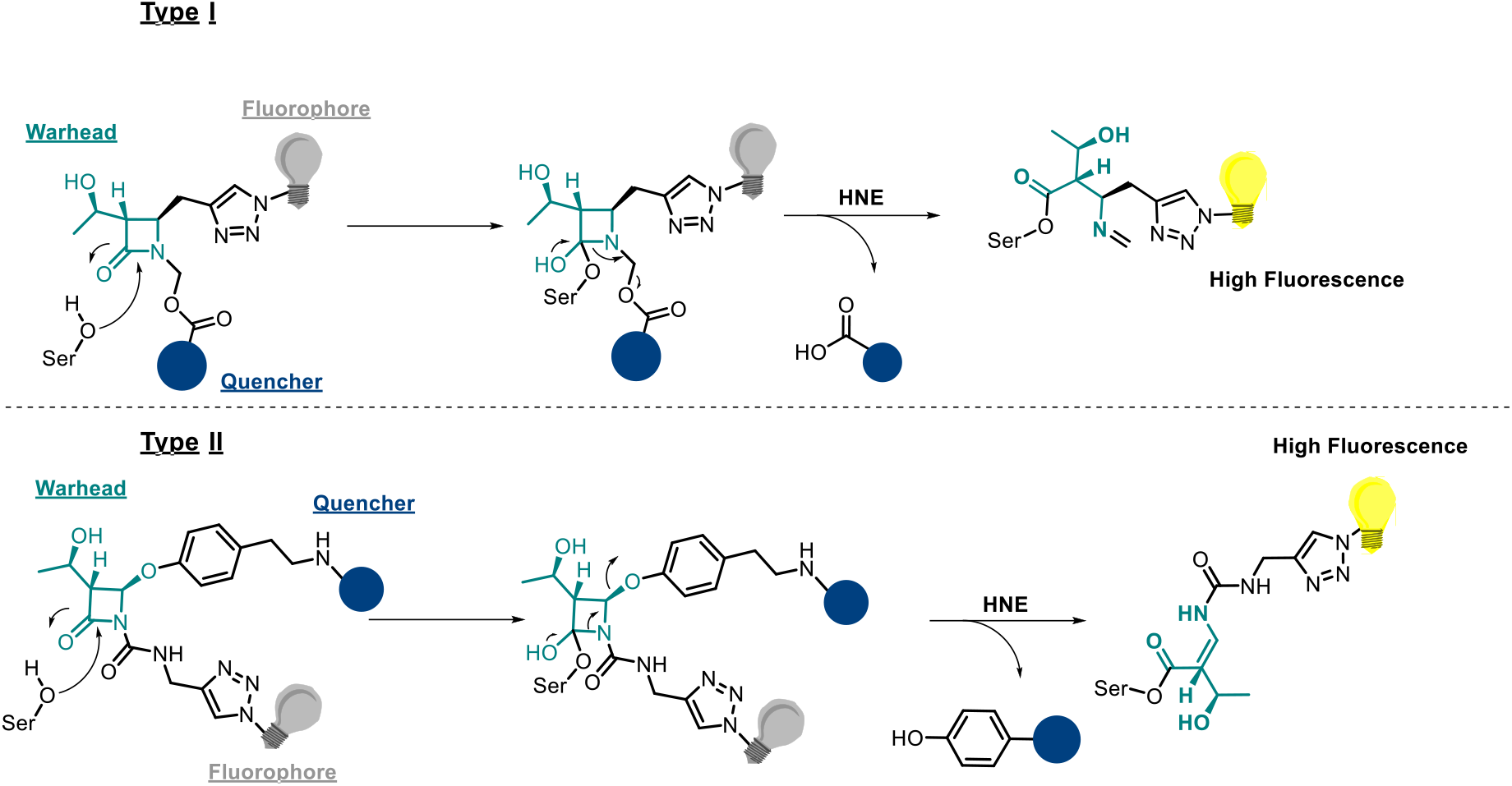
General qABP activation mechanism.

The qABPs incorporate the fluorophore BODIPY-FL and either a dinitrobenzene moiety or cAB40 as quenchers. For type I probes (Figure 4), synthesis began from the commercially available azetidine-2-one **1**. A Barbier-type reaction with propargyl bromide and zinc afforded the alkyne intermediate **2**, with retention of stereochemistry at the chiral centre, as confirmed by ^1^H NMR.^33^ Subsequent hydroxymethylation at N-1 and coupling of the quencher, yielded compounds **7i**-**iii** in good yields. Deprotection of the hydroxyethyl group using tetrabutylammonium fluoride, followed by the copper(I)-catalysed azide-alkyne cycloaddition with the previously synthesized BODIPY-FL azide **9**, furnished to the final type I qABPs **10**-**12**.

**Figure 4.**
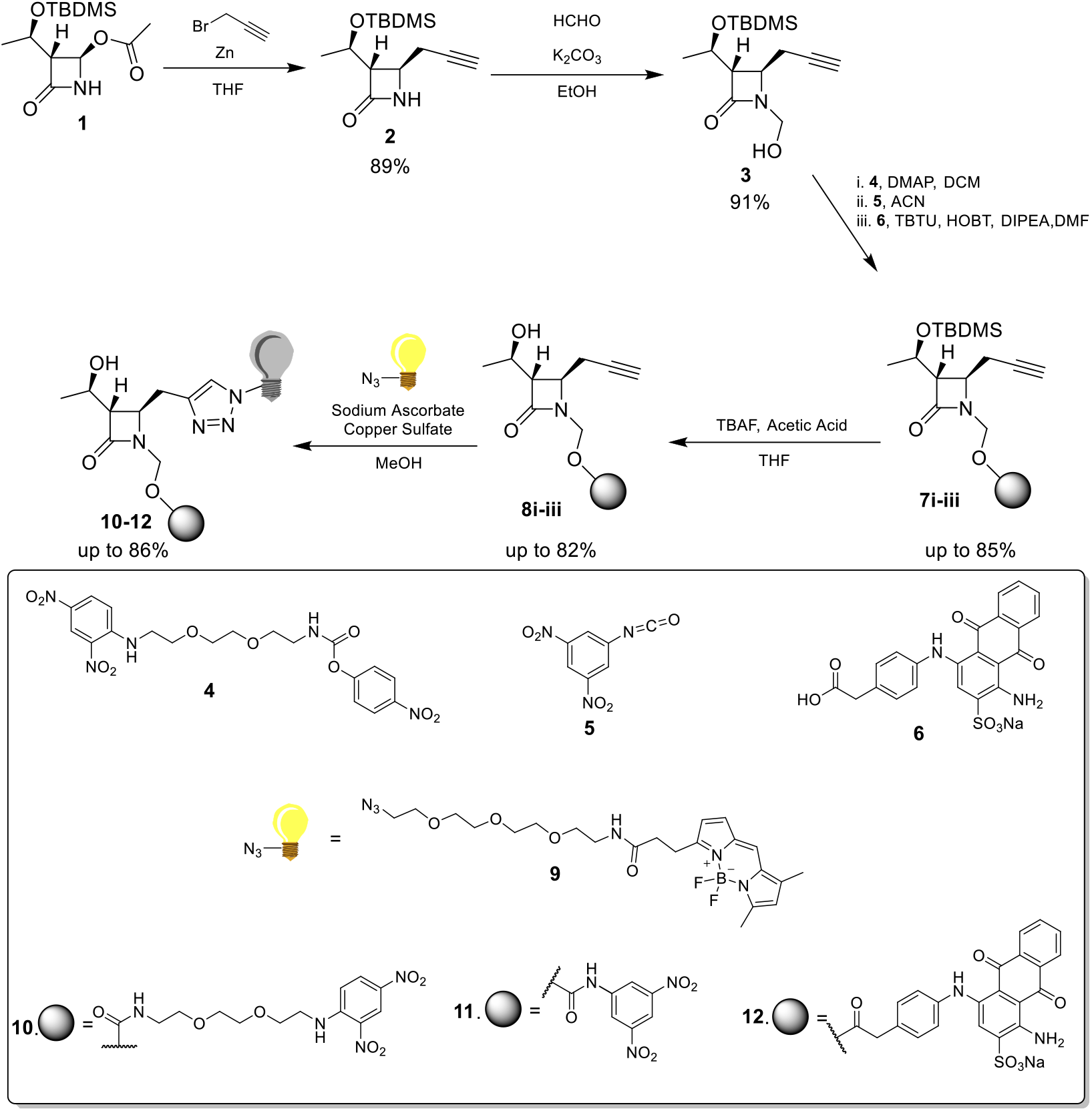
Synthetic strategy used to obtain type I qABPs. Compound **1** reacts with propargyl bromide through a Barbie-type reaction to achieve the enantiopure lactam **2**. Followed by a hydromethylation of the lactam ring to obtain compound **3**. The quencher moiety previously synthesised (**4**-**6**) was coupled with **3** generating intermediates **7i-iii** that was further deprotected (**8i-iii**) and finally a click reaction was performed to achieve the final qABPs (**10-12**).

The synthesis of type II probes began with azetidin-2-one **1**, followed by substitution of the acetoxy group with a phenoxy moiety bearing the quencher (Figure 5). Reaction with carbamate **17** afforded the alkyne intermediates **18i**–**iii**, which were subsequently deprotected. Final coupling with the BODIPY-FL azide via a copper(I)-catalysed click reaction yielded the type II probes **20**–**22**.

**Figure 5.**
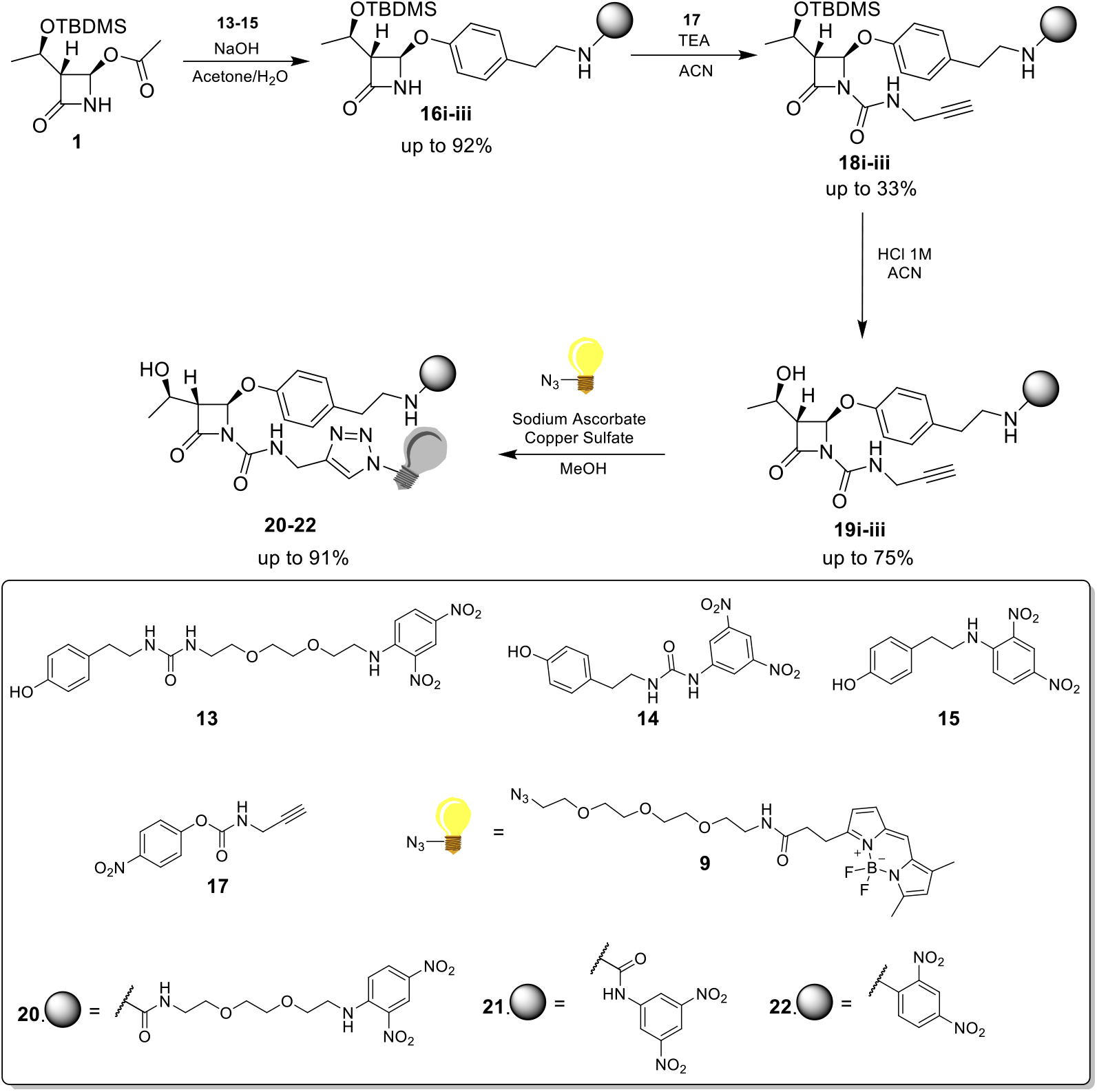
Synthetic strategy used to obtain type II qABPs. Compound **1** reacts with a phenol intermediate previously synthesised (**13-15**) that contains the quencher to obtain compounds **16i-iii**. Followed by a urea formation on the lactam ring to obtain compounds **18i-iii**. Compounds were deprotected (**19i-iii**) and finally a click reaction was performed to achieve the final qABPs (**20-22**).

### Photophysical properties

The photochemical properties were evaluated, and the stability of the probes was analysed in different pH values. All qABPs were stable in pH 7.4 PBS up to 48 hours. The alkaline hydrolysis of lactams was used as simple guide to determine the usefulness of the probes as enzyme acylating agents. To simulate the attack of the catalytic serine residue, the stability of the probes was assessed by UV-vis spectrophotometry in alkaline solutions. Inspection of the spectra revealed that type I probes where hydrolysed only at pH 13 (Figure S1). n contrast, their type II counterparts displayed a shift in the UV spectra at pH 10, consistent with *β*-lactam ring-opening and release of the quencher moiety. (Figure S2) To validate the ring-opening and consequently quencher released (turn-on of the probe) the reaction was monitored by fluorescence scan. For type I qABPs, the fluorescence increase persisted for up to 12 hours. In contrast, type II qABPs reacted more rapidly, achieving a comparable increase in fluorescence within 2 hours. This difference indicates that, in type II probes, quencher release is faster and more efficient, an essential feature for the intended application.

To evaluate enzyme activation, porcine pancreatic elastase (PPE) was used, given its high homology with HNE. The compounds were incubated with PPE, and fluorescence was measured over time. With type II qABPs, fluorescence increased up to sixfold within 3 hours (Figure 9); however, with type I qABPs, only a fourfold intensification was observed after 19 hours. These results showed that, in the case of type II, the reaction to release the extinguishing fraction was significantly faster compared to type I (Figure S3).

The fluorescence quantum yields (*ϕ*_F_) were calculated for all the compounds (Table 1) using an indirect method.^34^ A solution of fluorescein in 0.1M NaOH (*ϕ*_F_=89%) was used as a reference. While the azide used in probe synthesis had a fluorescence quantum yield of 100%, most probes synthesized in this work had a value lower than 11%, suggesting efficient quenching of the probe fluorescence. Probes **11** and **21** were the only exceptions, with a value higher than 20%, which could suggest that the amide bond to the dinitrobenzene moiety decreased quencher efficacy. After activation the fluorescence quantum yield increased up to 92%, validating our proposed turn-on mechanism and denoting that the compounds can be used as qABPs to detect the targets with low background fluorescence (Table 1).

**Table 1.** Fluorescence quantum yields before (PBS) and after activation (NaOH)

| | qABP | $\Phi_F$ (PBS) | $\Phi_F$ (After Activation) |
| --- | --- | --- | --- |
| Type I | **10** | 4% | 54% |
|  | **11** | 26% | 86% |
|  | **12** | 9% | 92% |
| Type II | **20** | 11% | 79% |
|  | **21** | 21% | 77% |
|  | **22** | 6% | 80% |

After the validation with PPE, we additionally validated probe **22** using pure HNE and human neutrophil lysate. These were incubated with the probe and the generation of fluorescence was measured through time. (<u>Figure 6</u>)

**Figure 6.**
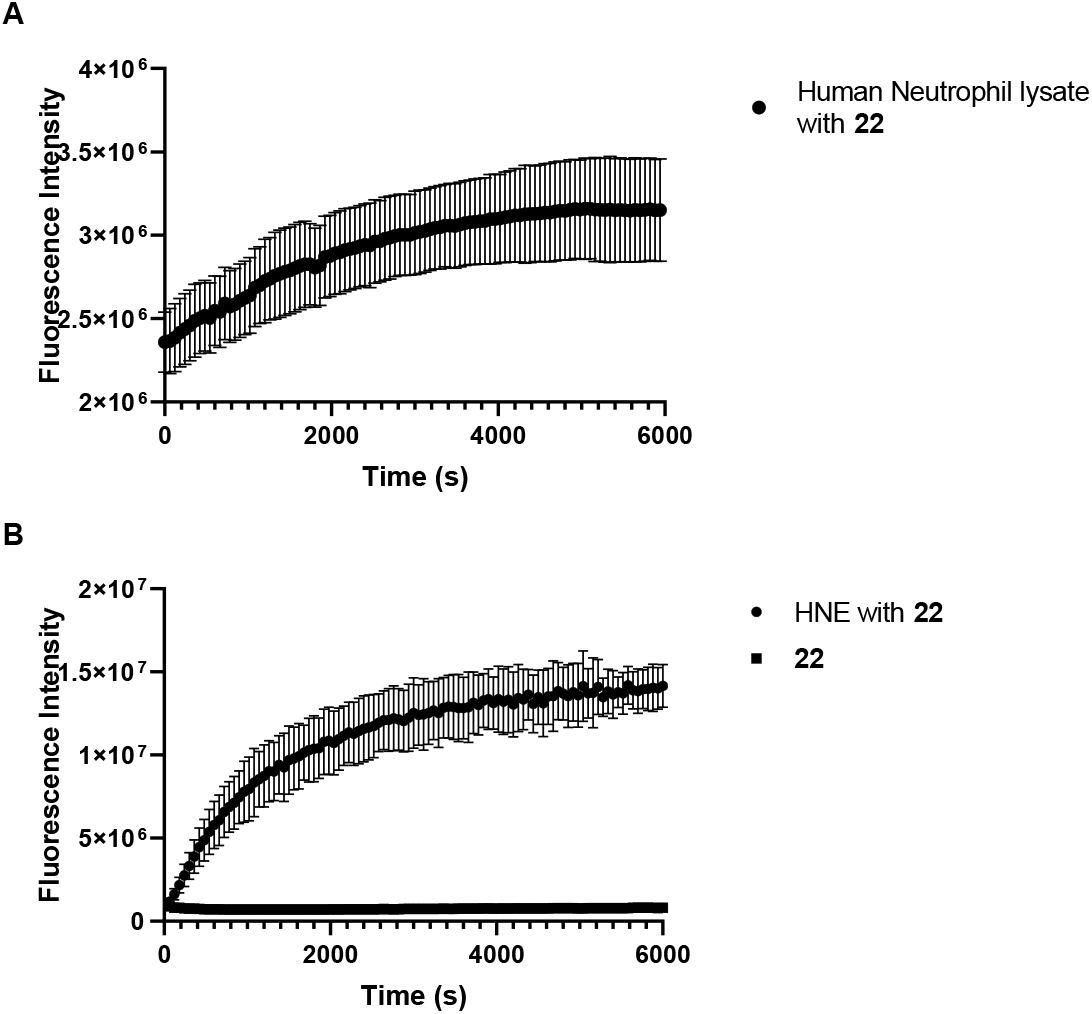
Left: Incubation of human neutrophils lysate with **22**. Right: Human neutrophil elastase with **22** and **22** in assay buffer. Fluorescence was measured every minute for 2 hours, using as excitation wavelength 485 nm and emission wavelength 535 nm. Assays were performed in triplicate.

In both cases, an increase in the fluorescence signal was observed (more significant in the case of pure HNE), confirming the reaction of the probe with the target and sequential release of the quencher moiety. The fluorescence of the probe in the buffer assay was also tested and remained very low and stable during the assay.

### Target Selectivity

Enzymatic assays were performed against HNE and other serine proteases including chymotrypsin, thrombin, kallikrein and urokinase. All probes displayed strong inhibitory activity against HNE, with IC_50_ values below 0.5 *μ*M. Type I qABPs **10** and **12** exhibited IC_50_ values of 0.50 and 0.12 *μ*M, respectively. However, due to their slower quencher release and delayed fluorescence response compared to type II probes, which limit their practical application, subsequent biological evaluations focused exclusively on Type II qABPs. In contrast, type II qABPs demonstrated high selectivity for HNE over related serine proteases, with IC_50_ values exceeding 10 *μ*M across the tested panel (Table 2). This selectivity is particularly noteworthy given the close similarity between PPE and HNE, which share approximately 43% sequence homology and highly similar active sites; consequently, PPE is often used in preliminary assays for HNE inhibitors.^35^

**Table 2.** Inhibitory potency of probes **20-22** against different serine proteases.

|  | IC<sub>50</sub> (μM) |  |  |
| --- | --- | --- | --- |
|  | 20 | 21 | 22 |
| **Chymotrypsin** | >10 | >10 | >10 |
| **Thrombin** | >10 | >10 | >10 |
| **Kallikrein** | >10 | >10 | >10 |
| **Urokinase** | >10 | >10 | >10 |
| **PPE** | >10 | >10 | >10 |
| **HNE** | 0.18 | 0.13 | 0.19 |

To further assess the selectivity of qABPs toward HNE in more complex environments, SDS-PAGE experiments were performed using both pure enzyme and HNE spiked into different proteomes. When incubated with purified HNE and analysed by gel electrophoresis, fluorescence was observed exclusively for Type II probes (Figure 7A). Notably, pre-incubation of the enzyme with ONO-6818, a potent and selective HNE inhibitor^36^, abolished the fluorescence signal (Figure 7B), indicating that the probes compete for binding at the HNE active site.

**Figure 7.**
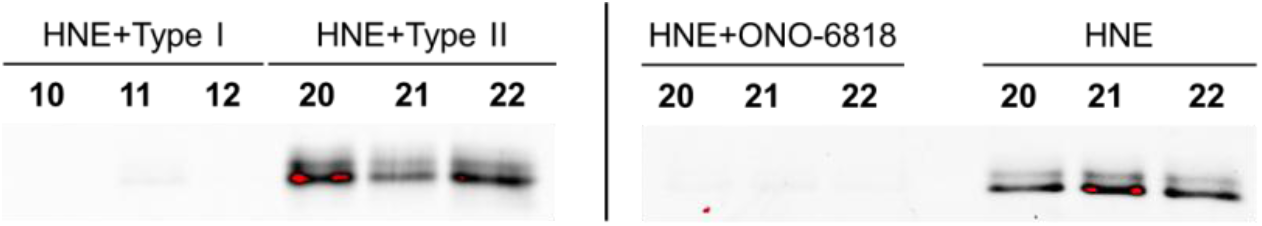
A) Incubation of the probes with HNE (left); B) Pre-incubation with ONO-6818 (right)

After confirming HNE engagement by gel, human embryonic kidney cell (HEK293) and epidermal carcinoma cell (A431) lysates were spiked with HNE and incubated with type II probes. The probes selectively detected HNE in the HEK293 proteome (Figure 8A). Probe **22** was also tested with decreasing concentrations of HNE, with minimal increase in background labelling and remaining selective for HNE down to 150 nM spiking (Figure 8B). Similarly, the A431 cell lysate experiment showed that all probes of type II could selectively detect HNE (Figure 8C), with probe **22** selectively detecting HNE down to a concentration of 75 nM of spiking (Figure 8D).

**Figure 8.**
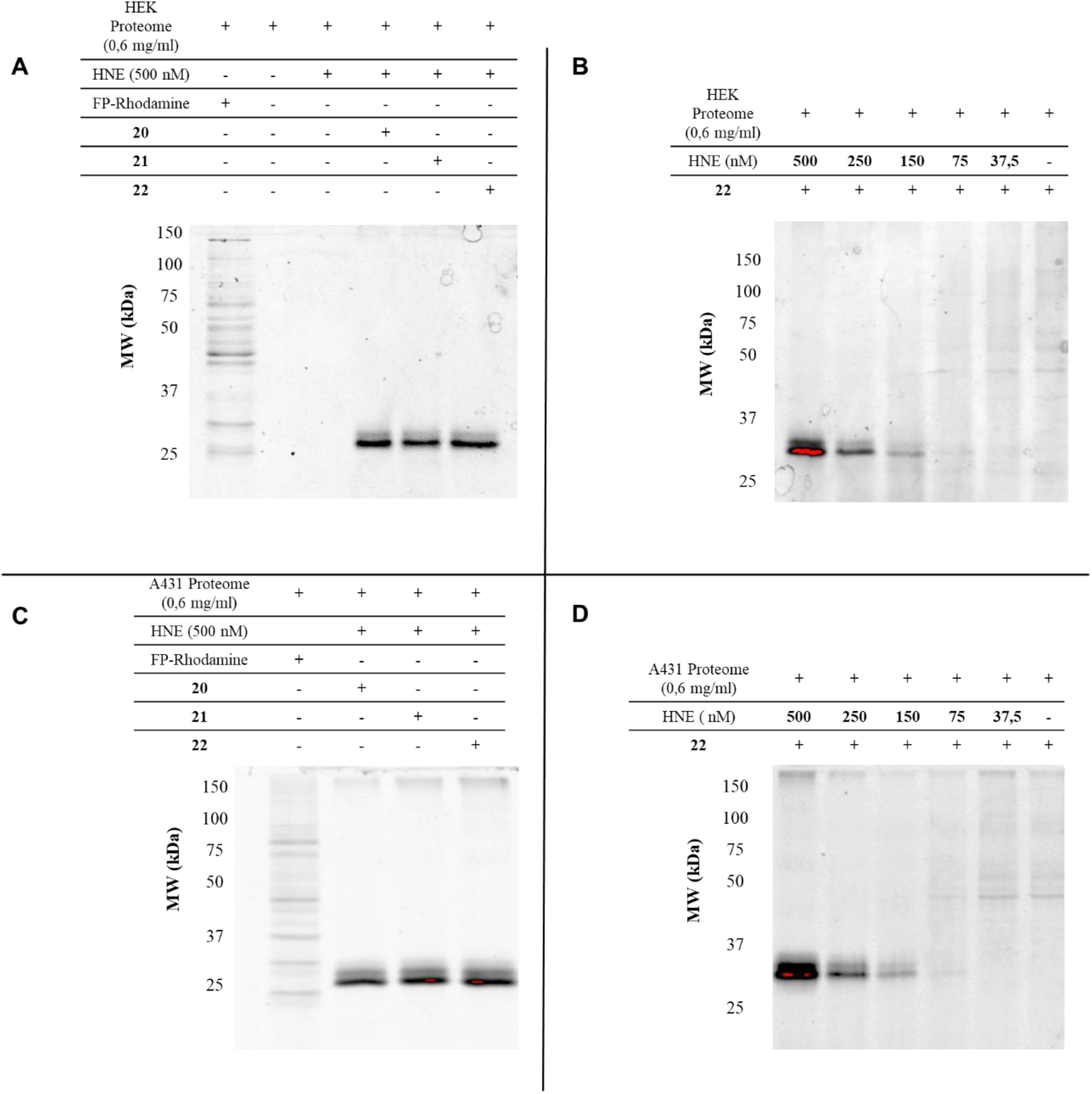
A) Type II probes incubated with HEK293 lysate spiked with HNE (500 nM); B) **22** incubated with HEK293 lysate spiked with decreasing concentrations of HNE; C) Type II probes incubated with A431 lysate spiked with HNE (500 nM); D) **22** incubated with A431 lysate spiked with decreasing concentrations of HNE.

Once the potential of the probes to target HNE was validated in a complex proteome, **22** was incubated with human neutrophils and U937 cell lysates (U937 is monocytic cell line that natively expresses HNE^32^). As presented in Figure 9, in both cases probe **22** was able to identify endogenous HNE in a complex proteome without requiring overexpression.

**Figure 9.**
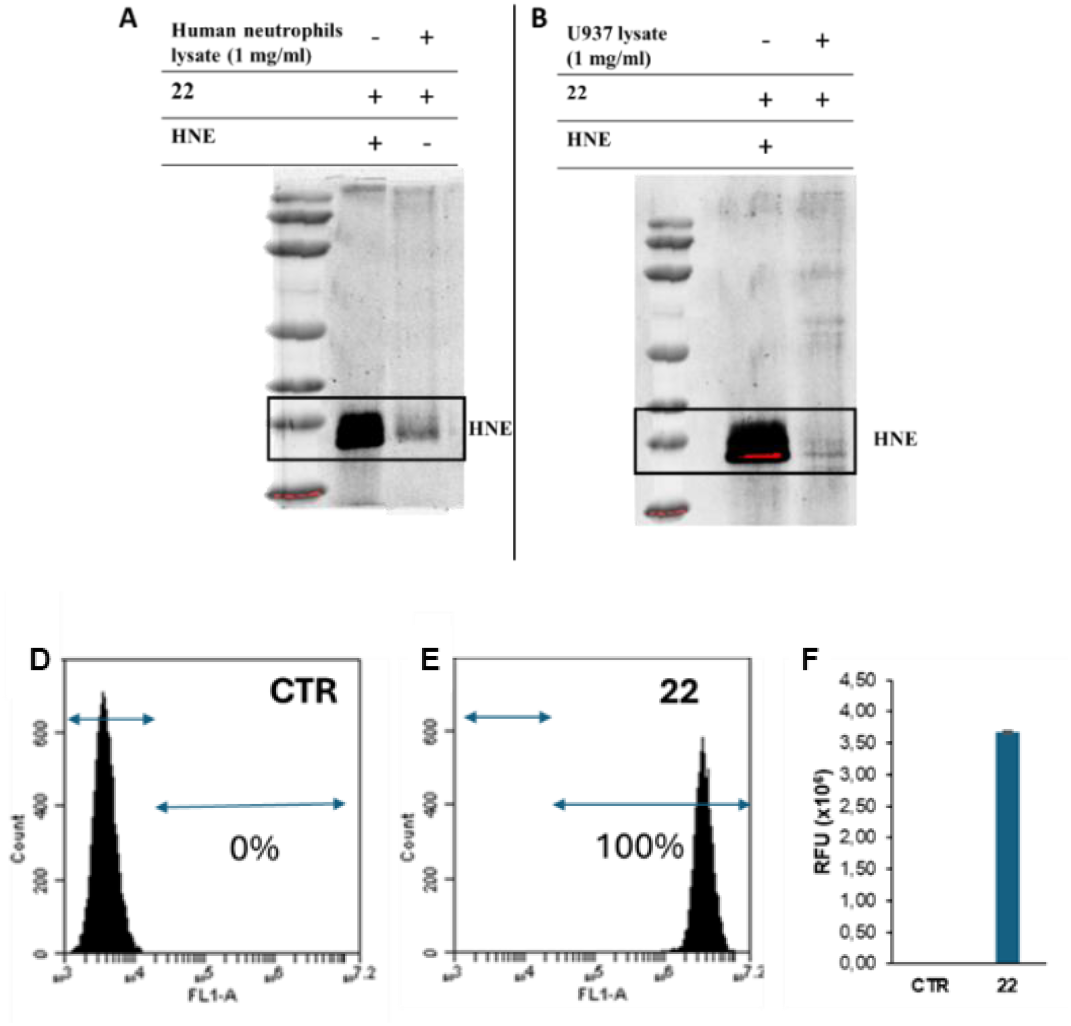
**A)** Probe **22** incubated with a lysate of human neutrophils, **B)** Probe **22** incubated with U937 cell lysate. **C)** Flow cytometry of living U937 cells. **D)** Flow cytometry of living U937 cells treated with **22** for 2h; in both cases the x-axis represents the fluorescence intensity measure, and the cell count is showed in the y-axis. **E)**. Bars graph (Mean+SD) represents the median fluorescence intensity of the control and the cells treated with **22**.

Encouraged by these promising results, we next evaluated whether the synthesised probes could be internalized by living cells. Cellular uptake was assessed in U937 cells and human neutrophils using flow cytometry and fluorescence imaging, respectively. In U937 cells, flow cytometry analysis showed that control cells (untreated with **22**) exhibited no detectable fluorescence, whereas cells treated with **22** displayed a marked increase in fluorescence intensity, indicating efficient probe internalization and activation (Figure 9).

For human neutrophils, internalisation was assessed by fluorescence imaging using a high-content analysis system. A strong fluorescence signal was observed in the cytoplasm following incubation with **22**, whereas the negative control showed no detectable fluorescence (Figure 10), further confirming efficient cellular uptake.

**Figure 10.**
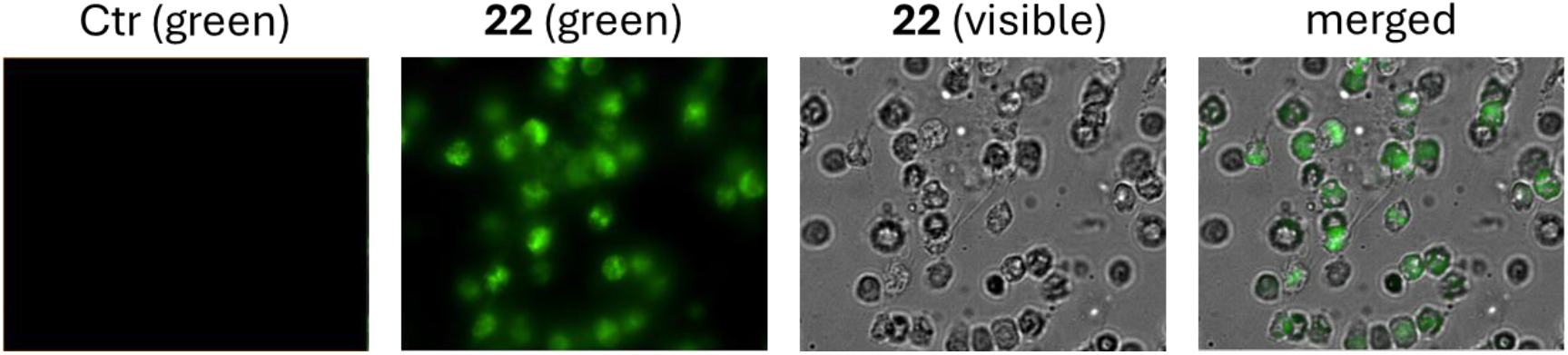
Incubation of **22** with human neutrophils. Images were taken after 2h of **22** additions.

Overall, the BODIPY-tagged *β*-lactam probes presented remarkable selectivity and strong potential as qABPs for detection of HNE in complex proteomes, highlighting their value as tools to elucidate the role of HNE in the tumour microenvironment. To further investigate the structural features underlying this high selectivity, both covalent and non-covalent docking studies were performed using compound **22**.

Molecular docking of probe **22** showed that the hydroxyethyl substituent at C-3 of the *β*-lactam warhead occupies the primary binding S1 pocket, with the hydroxyl group forming a hydrogen bond with Val216 (Figure 11A). This observation suggests that the predominantly hydrophobic S1 pocket can also accommodate polar interactions.

**Figure 11.**
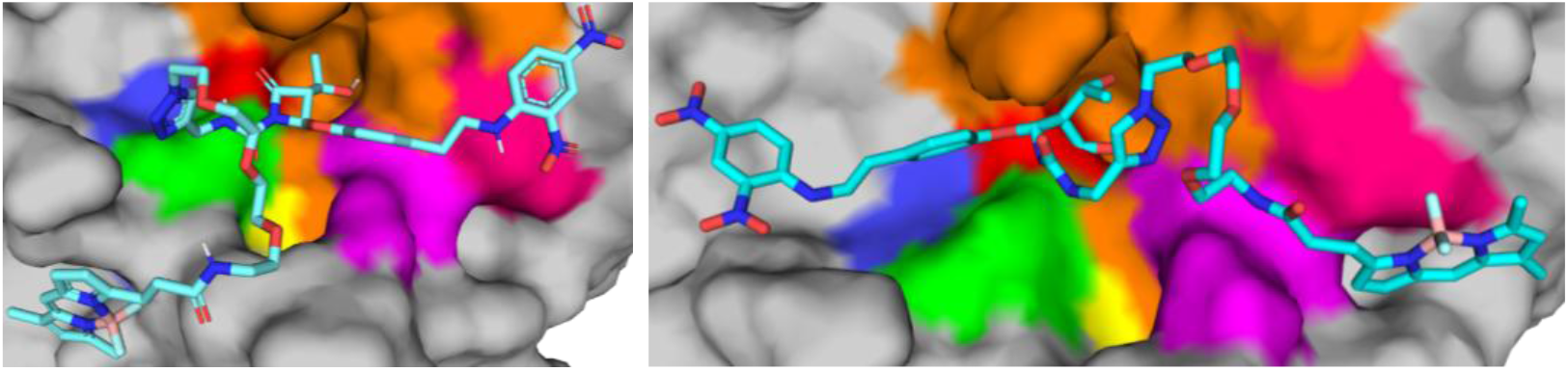
A) Non-covalent docking pose (right) of **22** in the active site of HNE, obtained from PDB 9SI4. Active site is highlighted with different colors, Ser-195 in red, His-57 in green and Asp-102 in yellow. B) Covalent docking pose (left) of **22** in the active site of HNE, obtained from PDB 9SI4. qABP **22** is presented as blue sticks and PDB 9SI4 is presented as a grey surface. Active site is highlighted with different colors, Ser-195 in red, His-57 in green and Asp-102 in yellow. Pictures were prepared using Pymol. (S1 pocket represented in orange, S2 pocket represented in pink, S1’ represented in blue, S3 represented in hotpink).

Regarding covalent docking (Figure 11B), the results indicate that probe **22** establishes multiple interactions within the S1 pocket of HNE, primarily involving residues Phe41, Phe192, Gly193, Ser214, Phe215, Val216, and Arg217. Additional interactions were also observed at the enzyme surface, mainly with Leu35, Leu167, Ser214 (S1) and Val216 (S3). These findings support the capacity of the S1 pocket of HNE to accommodate polar substituents.

Overall, the synthesised probes presented remarkable selectivity and a strong potential to be used as qABPs to detect HNE in complex proteomes and could become an important tool to reveal the roles of HNE in the tumour microenvironment.

## Conclusion

In this work, we describe a synthetic strategy to generate qABPs capable of selectively targeting HNE in biological media and complex proteomes without the need for a peptidyl linker for molecular recognition. The developed qABPs exhibited high quenching efficiency, with FQY values below 20%, which increased to up to 92% upon activation. Probe activation was confirmed through fluorescence assays using both purified HNE and human neutrophil lysates.

The probes displayed remarkable selectivity for HNE, even when compared to closely related serine hydrolases. Their effective detection and activation in biological matrices were further demonstrated across three different cell lines, with and without HNE spiking, including human neutrophil lysates.

Overall, this study provides a set of versatile chemical tools for investigating the role of HNE in the tumour microenvironment and supports its validation as a relevant target in cancer therapy.

## Author contributions

RM and RF conceived and designed the study. Synthesis was developed by RF and AM-A. Photophysical and biology data collection, and data analysis were carried out by RF, LC, AMM and LG. Flow cytometry and cell imaging studies were performed and analysed by MCR, RFG and OG. Computational studies were performed by IR and RG. All authors analysed the results and contributed to writing the manuscript.

## Acknowledgements

This project has received funding from the European Union’s Horizon Europe Programme under grant agreement No 101132028 (project IMPULSE), and from Fundação para a Ciência e Tecnologia (FCT) trough projects 2022.07857.PTDC and UID/04138/2025 (doi: 10.54499/UID/04138/2025), UID/PRR/04138/2025 (doi: 10.54499/UID/PRR/04138/2025), UID/PRR2/04138/2025 (doi: 10.54499/UID/PRR2/04138/2025). FCT also funded fellowships SFRH/BD/137459/2018 (RF) and COVID/BD/153228/2023 (RF). We also acknowledge support from ERIC EU-OPENSCREEN, PT-OPENSCREEN and National NMR Network, supported by Infrastructure Project Nº02216, co-financed by FEDER (COMPETE2020, POCI, PORL) and FCT (PIDDAC), and Portuguese MS Network, LISBOA-01-0145-FEDER-022125, supported by Lisboa2020, under the Portugal2020 Partnership Agreement, through the European Regional Development Fund. The bioimaging system was acquired by Fundación MEDINA through an Andalusian Government (Spain) infrastructure and equipment grant under the PAIDI 2020 and RIS3 Andalucía strategies (reference: IE_18_0031).

